# Radical pair based magnetic field effects in cells: the importance of photoexcitation conditions and single cell measurements

**DOI:** 10.1101/2022.11.09.515724

**Authors:** Jonathan R. Woodward, Noboru Ikeya

## Abstract

A recent publication^1^ on the bioRxiv preprint server aims to replicate our observation of magnetic field effects on the autofluorescence of HeLa cells^2^, but is unable to reproduce the effects described in our original work. Here we examine this new study and demonstrate, based on a model of the reaction photocycle, why the differences in the measurement conditions used render the experiment unlikely to be able to observe the originally reported effect. In addition, we highlight substantial problems in the quality of the data in the replicate study and reiterate the advantages of the direct modulation, single cell measurement approach presented in the original work over a more standard statistical approach.

## Introduction

The observation of magnetic field effects (MFEs) in general, and specifically those attributed to the radical pair mechanism (RPM) has had a chequered past. For example, measurements of MFEs on enzyme-based reactions (e.g. refs^3-5^) have proven difficult to replicate^6-8^, and indeed one of the authors of this manuscript has co-authored past studies which failed to replicate previously reported effects^6-7^. More recently, the issue is particularly pertinent to studies of MFEs on reactive oxygen species (ROS) in biology, which has led to uncertainty over their robustness. Therefore, the replication of MFE observations in biology is critical and necessary, but suffers from the broader issue of the reluctance of scientific journals to publish replication studies and the obsession in modern science of novelty over reproducibility.

We were thus delighted to recently encounter a pre-print that aimed to replicate our original study of magnetic field effects on the autofluorescence of HeLa cells (although disappointed that the authors at no point contacted us to discuss details of our work before publishing the preprint). While we worked very hard in our original study to minimize all conceivable possibilities of errors, we had hoped that other groups would work to reproduce our findings and validate our observations or identify inconsistencies. In particular, we were impressed that the scientists involved in replicating our study had worked hard to try to match the conditions of our original experiment. Initially surprising for us was the fact that the group were unable to reproduce our original findings and went on to take a different, more conventional approach based on statistical measurements. The latter study also failed to find any observable effects of magnetic fields of up to 500mT on the autofluorescence of HeLa cells. In order to explain the discrepancy between the results, the authors were quite critical of our original methodology and suggested that our analysis method was unclear and did not employ appropriate statistical considerations. In this paper we would like to address the specific technical reasons why we believe the replicate study failed to reproduce our observations (and indeed as performed is not capable of ever reproducing our observations), to draw attention to the problematic data sets presented, and finally to rebut the criticisms made about our real-time, modulation based single cell measurement approach. Indeed, we believe that this approach is what made our original experiment compelling and where possible should be a model for other scientists trying to provide conclusive evidence of MFEs in biology.

### Measurement conditions in our original work and the replicate study

To a large extent, the study by Uzhytchak *et al*. reproduces the experimental and instrumental conditions of our original study effectively. However, there is one aspect where the measurements differ substantially, and unfortunately this is perhaps the most critical for the observation of magnetic field effects. The difference is in the photoexcitation conditions.

In our original work, the microscope was configured to strike a careful balance between the size of the irradiation region (the region of interest, ROI) and the laser power. We took a 450nm diode laser (the optimal visible wavelength for flavin photoexcitation) and cleaned the beam using a single mode optical fiber. The output of this beam was then nearly collimated before entering a high (1.49) NA objective lens. The size of the collimated beam deliberately did not fill the back aperture of the objective and the beam divergence was carefully controlled to produce a spot of diameter 12 μm at the sample. This size was selected, based on the average size of HeLa cells, to allow irradiation of a substantial subcellular ROI, without irradiation of nearby organelles (particularly the nucleus), or of neighbouring cells. Thus, our measurements were performed not with a confocal arrangement (as typical commercial laser scanning or spinning disc microscopes employ), nor with an epi-arrangement, irradiating a larger region of the sample (with a necessary reduction in laser intensity). In addition, it means that we can image the sample in the region of interest using continuous or pseudo-continuous irradiation and record in real time using the sCMOS camera, with precise control over the power to control the photocycle kinetics. The results of this are clear in the movie supplied in the supporting information of our original article^2^. Our laser was operated at a 50% duty cycle at a frequency of 100Hz. We experimented extensively with the laser power, duty cycle and switching rate based on existing literature^9^ to strike the correct balance between sufficient power in the on cycle (to optimise photocycle kinetics, see later) and the reduction of photobleaching. The conditions selected represent the optimum conditions for resolving the magnetic field effects. Thus our conditions can be considered as continuous irradiation intensity of 1 kWcm^-2^ for 5ms and then laser off for 5ms (giving an average irradiation intensity of 0.5 kWcm^-2^).

In the replicate study by Uzhytchak *et al*., the photoexcitation conditions are significantly different. First the study uses a different excitation wavelength of 488 nm (still effective for flavin photoexcitation although with an absorption cross section reduced by about 30 %^10^). Much more critically, the study uses a spinning-disc confocal arrangement. To confirm the details of the system used by the original authors, we consulted the specifications published by the manufacturer (Yokogawa) of the instrument used (CSU-W1) and confirmed some specifications with the manufacturer directly. In such a microscope, the excitation beam is focussed into pinholes through a series of microlenses, resulting in a series of discrete locations being irradiated simultaneously (**Fig.1 a**). In this particular instrument, the disc contains a total of about 20000 microlenses / pinholes, with about 1000 irradiated at any one time^11-12^ (**Fig.1 b**). As the disc spins, the irradiation positions move (based on the pinhole pattern in the disc) and the pattern repeats 3 times each rotation based on the fact that this model uses 3 spirals (1/3^rd^ of a rotation). This means that any particular point in the sample experiences short periods of irradiation as a beam passes over it. This is equivalent to pulsed photoexcitation at any particular point in the sample, with each location in the sample being briefly photoexcited and then returned to the dark until another beam passes that position or the original beam returns after a 120 degree rotation of the disc. According to the manufacturer’s specifications (see this tutorial^13^ for an explanation), the spinning disc unit used has a minimum possible spinning speed of 2000rpm (to match the camera exposure time with the disk speed and avoid streaking), when operated with a camera exposure time of 100ms (as the authors employed). The maximum possible spinning speed is 4000rpm. These spin speeds mean that the entire field of view is completely scanned at a repetition rate of 100Hz or 200Hz (2000 or 4000rpm x 3 image scans per revolution / 60 seconds per minute). For our calculations, we will assume the slower repetition rate, which will favour stronger MFEs.

**Figure 1.**
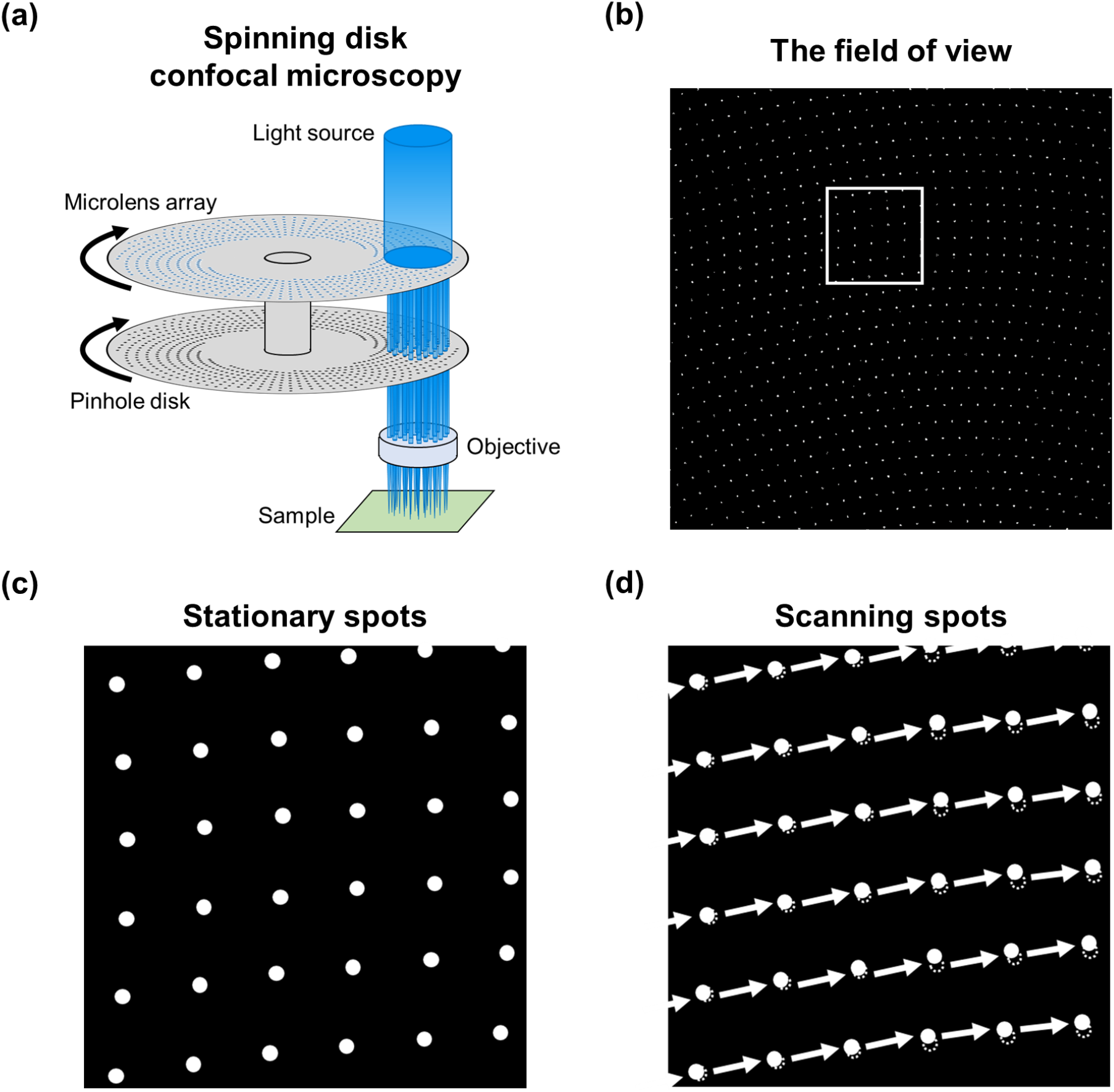
Principle of a spinning disc confocal microscopy. (a) Schematic of the illumination system (adapted from ref^22^.). (b) An example of the field of view with a spinning disk confocal unit (Yokogawa CSU-W1) (adapted from ref^23^.) (c) The stationary beam spots (zoomed in (b)). (d) The scanning beam spots.

It is important to know what laser power reaches each point in the sample as a beam passes over. According to the authors, the laser power was around 3mW (2.95mW). The authors claim an overall irradiation intensity of 0.4kWcm^-2^, but this cannot correspond to the average power across the entire sample (~130μm x 130μm [shown in their Movie S1]) as this could be a maximum of 0.0175kWcm^-2^ shown in the following calculation.

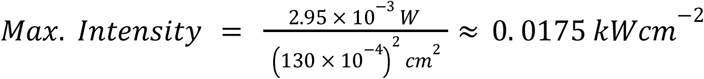

Therefore, we can assume that the intensity refers to the beams delivered through each microlens in the spinning disk array. Given that there are 1000 beams hitting the sample at any given time, the estimated irradiation spot diameter of each beam can be estimated as 969 nm as follows.

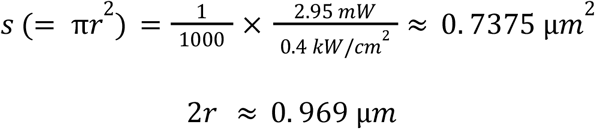

Thus, about 1000 beam spots of the above diameter exist in the field of view at any one time, move as the disk rotates, and scan the entire field of view (as shown in **Fig.1 c, d**). For the 2000 rpm case, it takes 10 ms to complete one scan of the entire field of view. Since the movement of the spot is driven by the rotation of the disk, the spots in the field of view move at different speeds on the inner and outer circumference, and unless the spinning disk pattern is accurately known, we are not able to fully know the photoexcitation profile at each point during a single scan. Here, we estimate the irradiation conditions of the replicate study by determining average values.

If the spinning speed is 2000 rpm, 10 ms is required to scan the entire field of view. Therefore, the scanning speed Δ*S* is as follows.

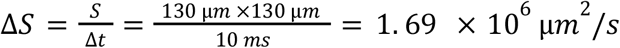

Since about 1000 spots always scan the field of view at any given time, the scanning speed per spot Δ*s* can be estimated as follows.

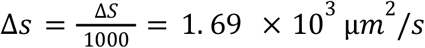

And, the irradiation time of one spot is estimated as follows

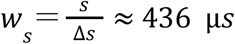

However, this value is an estimate in which the entire field of view is scanned without any overlapping of scanning orbits drawn by the spots. In actual spinning disk microscopy, each point (pixel element) in the field of view is scanned multiple times because there are overlaps in the trajectories scanned by the individual spots. In other words, the scanned area is larger than the entire area of the field of view. Taking this into account, we next estimate the irradiation time for each point and the number of times it is irradiated.

Let *S*’ be the total area scanned by the spinning spots during one scan. Then the average number of excitations of all pixels during the formation of one frame can be defined as follows.

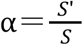

Therefore, the scan area speed per spot can be written as follows.

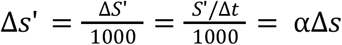

Then, the irradiation time per spot at a given point can be expressed as follows.

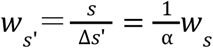

Therefore, if we estimate the total area scanned by the spots, we can find the average value of the irradiation time and the number of irradiations at each point. Here, we estimate the average value of the maximum total area scanned by the spinning spot.

First, the spinning disk has approximately 20000 microlenses and pinholes in total, and one image is formed by 1/3rd of a rotation, which means that one image is formed by 20000/3 different spots passing through the field of view. Assuming that all spots pass in a straight line horizontally for simplicity, we can estimate the maximum of the total area scanned by the spots.

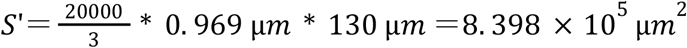

and the average number of excitations at any pixel is estimated as follows.

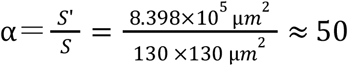

The average of the maximum irradiation time at any point is then estimated as follows.

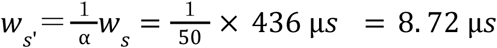

Therefore we can conclude that a reasonable description of the photoirradiation conditions of any particular point in the sample is equivalent to a pulsed irradiation source of wavelength 488 nm with a peak power of 0.4 kWcm^-2^ and a pulse length of 8.7 μs operating at about 5 kHz. In practice most pulses will be shorter than this and there will be periods during each cycle where the repetition rate is lower or higher and these will vary with the position in the image. Contrast this with the irradiation condition in our original experiment of a pulsed irradiation source of wavelength 450 nm with a peak power of 1.0 kWcm^-2^ and a pulse length of 5 ms operating at 100 Hz. These two irradiation schemes are presented in **Fig 2a**.

**Figure 2.**
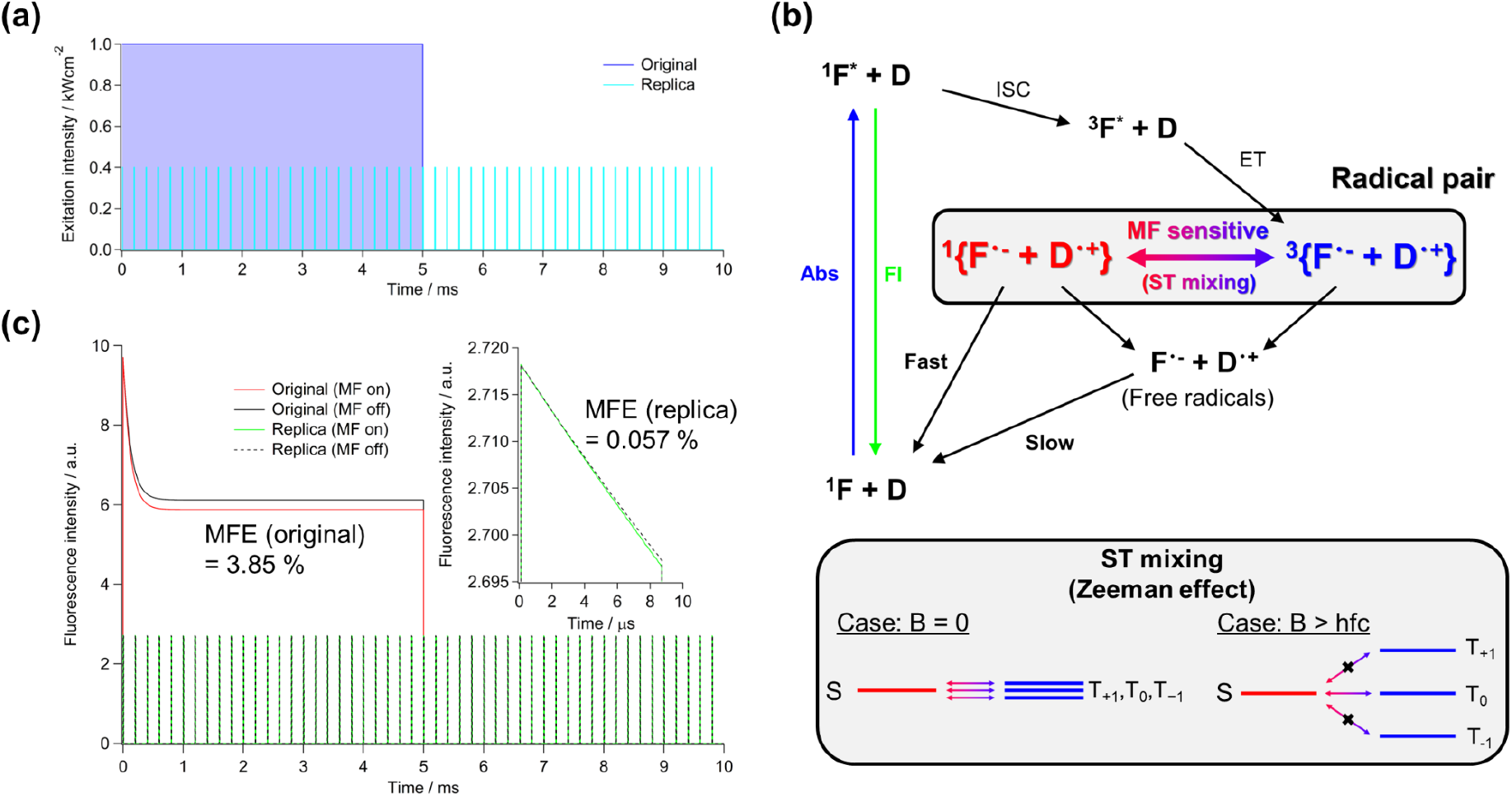
Simulated magnetic field effects with irradiation conditions from the original and replicated studies. The following rate coefficients were used in the simulation: laser excitation, 2.0 × 10^4^ s^-1^ (original), 5.6 × 10^3^ s^-1^ (reproduction), sum of fluorescence and internal conversion rate coefficients, 1.7 × 10^9^ s^-1^, radical pair formation, 3.6 × 10^8^ s^-1^, back reaction to flavin ground state from singlet RP, 1 × 10^5^ s^-1^, RP singlet-triplet (ST) state mixing for all triplet states (T_+1_, T_0_, T_-1_) (zero field) and only T_0_ (high field), 8.0 × 10^7^ s^-1^, formation of free radicals, 1 × 10^5^ s^-1^, slow return of free radicals to flavin ground state, 5 × 10^3^ s^-1^. (a) Irradiation schemes of the original and replica study. (b) Flavin based photocyclic kinetic model. For calculating the magnetic field effect, the Zeeman effect on the triplet states is employed. In zero field, the singlet (S) state can mix with all three triplet (T) states (T_+1_, T_0_, T_-1_), while in an applied field greater than the average hyperfine couplings (hfc) in the RP, the T_+1_ and T_-1_ states are separated in energy and cannot undergo mixing with the S state. (c) Comparison of the simulated magnetic field effects on fluorescence for the original and replica studies. MFE = (I(on) -I(off))/I(off). The MFE is calculated over the time period of 100 ms (one camera frame).

For this discussion to be relevant to the different observations in the two studies, it is important to understand why the excitation intensity and duration is critical. Indeed for the vast majority of flash photolysis studies of MFEs on chemical reactions proceeding through RP intermediates, the excitation intensity does not influence the magnitude of the MFE, but only the signal-to-noise ratio in the measurement, based on the number of molecules excited. A sufficiently strong excitation pulse is necessary only to generate enough RPs (typically observed using transient optical absorption) to resolve changes of a few percent. In the case of cyclic flavin photochemistry, however, the excitation laser has two roles. The first is to photoexcite the sample and drive the photochemical generation of radical pairs, and the second is to monitor the return to the oxidised flavin ground state (based on the fluorescence it emits when photoexcited) to report on the fraction of RPs quickly undergoing back electron transfer through the spin-selective singlet channel. The result of this is that continuous irradiation is used to drive the photocycle into equilibrium and develop magnetic field sensitivity in the ground state flavin concentration. As a result, the magnitude of the observed MFE becomes significantly dependent on both the continuous irradiation power and the time for which the irradiation is applied (i.e. saturating the MFE at equilibrium takes time). This can be readily observed in the laser power dependence of model flavin photoreactions – for example aqueous solutions of flavin adenine dinucleotide. Too little or too much laser power results in a reduction of the observed magnetic field effect. Our original study was designed based on this understanding, and early experiments focussed on establishing the optimum irradiation conditions. Increasing or reducing the laser power in our measurement too much resulted in difficulties in observing a MFE.

To explain this issue more clearly, consider the flavin photocycle kinetic model presented in **Fig. 2b** (which is based on the scheme presented in our original study). Flavins are photoexcited to the singlet excited state from which they can return to the ground state radiatively (the observed autofluorescence) or non-radiatively, or they can undergo intersystem crossing (ISC) and the resulting excited triplet state can undergo electron transfer (ET) from a donor molecule to generate a triplet-born RP. The triplet-RP can either generate uncorrelated free radicals or undergo coherent spin-state mixing to generate the singlet-RP. The singlet RP can either generate uncorrelated free-radicals (FR), undergo spin-selective back electron transfer to the ground state or undergo coherent spin-state mixing to regenerate the triplet RP.

For low excitation rates, the RP / FR concentrations established are very low and so the changes in the ground state concentration of flavin are a very small fraction of the total concentration. When the excitation rate becomes larger than the slow return to the ground state (which is not spin-selective), then the RP / FR concentration increases and the ground state flavin concentration is reduced. Under these conditions, the rate of repopulation of the ground state through the spin-selective reaction is significant and the ground state population becomes magnetic field sensitive. If the excitation rate increases sufficiently to outpace the rate of spin-selective reaction, the ground state flavin concentration becomes very small and the fluorescence signal difficult to measure. Furthermore, if photoexcited flavin molecules can also undergo a non-reversible alternative reaction (corresponding to photobleaching), then at high laser powers, flavins are rapidly recycled and efficiently photobleached.

Understanding the effects of the photocycle and photobleaching makes it clear that finding the optimum photoexcitation conditions for the measurements is critical, particularly in the case of cellular measurements where the sample cannot be replaced through diffusion of fresh molecules. It also highlights the need for continuous irradiation to establish equilibrium in the photocycle and render the fluorescence signal sensitive to the RP dynamics. A flash photolysis experiment (with e.g. nanosecond laser flashes) measuring flavin fluorescence cannot observe MFEs, as the fluorescence arises from the ground state flavin concentration before the RP reaction has occurred. Only if the repetition rate of the laser occurs such that the time between pulses is on the same timescale of the FR or shorter, will MFEs be measurable. The estimated time between same spot irradiations in the replicate study is around 200 μs, by which time almost all RPs will have returned to the ground state or possibly moved from the irradiation region (depending on their mobility within the cell - this is not the case in our large continuous irradiation beam which is also not depth resolved).

To provide a much more robust demonstration of why the reproduction study is incapable of observing the magnetic field response, we performed kinetic simulations, based on the reaction scheme in **Fig. 2b** for the irradiation conditions used in the two experiments. We used photophysical parameters based on known flavin chemistry^13-14^ and kinetic parameters suggested for the cryptochrome reaction cycle^15^ to give as realistic a model calculation as possible (parameters are provided in the figure caption). The rate coefficients were the same for both experiments, with the exception of the excitation rate coefficient, which was multiplied by a factor of 0.4*0.7 for the reproduction study, to compensate for the reduced laser intensity and change of absorption cross section at 488 nm. **Fig 2c**. shows the results of the simulations. The calculated total MFE on fluorescence is **3.85%** for the original irradiation conditions and **0.057%** for the reproduction study. Examination of the time dependence shows why this is the case. The light source needs to be applied for long enough to establish equilibrium in the reaction cycle and the spinning disc configuration moves the irradiation spots too quickly for this to be possible. The result is a dramatic difference in the sensitivity of the observed fluorescence to the MFE.

The irradiation conditions in the replicate study cause other practical problems. In our original measurements, the signal-to-noise ratio of the fluorescence signal is high (>4000 at time of weakest signal). It is this that renders our measurements easily able to resolve magnetic field effects of less than 1%. Uzhytchak *et al*. identify the problem of Poisson noise in their images, but our signals are much stronger due to the much higher average excitation intensity and thus amount of integrated fluorescence (the kinetic simulations suggest 26 times greater integrated fluorescence intensity in our measurement), along with our greater fluorescence capture efficiency (primarily due to greater objective lens NA). This means that Poisson noise is negligible relative to other sources of noise. A comparison of our video with those in the replicate study highlights the enormous signal-to-noise advantage of our measurements, with clear camera noise observable at all times in the latter.

In summary then, the irradiation conditions generated by a spinning-disc confocal arrangement are incompatible with the flavin photocyclic reaction scheme and would not be expected to lead to observable MFEs on the autofluorescence signal. We would like to suggest that the authors change the irradiation conditions to closely match those in our study in order to correctly evaluate our findings.

In addition, it is worth highlighting the fact that we provided a positive control measurement based on an established magnetic field effect on a chemical system (flavin adenine dinucleotide in pH 7.4 buffer at an endogenous flavin concentration level (~ μM)^16^ with a detection volume at cell volume level (~pL)^17^). Our optimised irradiation conditions allow clear observations of the weak MFE in this system while minimizing photobleaching of the flavin^9^. It would only be reasonable for the authors of the replicate study to suggest that our measurements are problematic if they could first demonstrate that their own measurement conditions are capable of reproducing the observed MFE in this system (more specifically, at endogenous flavin concentration levels and cell volume levels). Such a positive control confirmation of the experimental conditions and microscope sensitivity is conspicuous by its absence and is likely indicative of the inappropriate irradiation conditions employed.

### Specific criticisms of our data analysis

The report of the replicate study criticizes both our measurement approach and our data analysis. Here we address these criticisms, beginning with the latter.

Uzhytchak *et al*. state, *“In the original study [25], it is noted that cells showed a magnetic response with a magnitude of 1 to 2*.*5%. These are very weak changes. Guidance for quantitative fluorescence microscopy state that for a change of even as high as 25% between two conditions, sampling of ~ 100 cells for each condition is required to measure the change with statistical confidence [42]*.*”* This statement alone clearly highlights that the authors do not understand the difference between a statistical study of a measurement group vs a control group and our study in which no control group is necessary because the magnetic field imparts a real time change in flavin fluorescence on an individual cell (i.e. that the same region of the same cell serves as sample and control). We fully agree that 1 - 2.5% changes are always going to be challenging for statistical biological assays and that this has led to countless problems in the field in the past. In our experiments, we can capture the fluorescence signal with sufficient signal-to-noise ratio that if a correlated change in the fluorescence occurs with the application of an external modulation (in this case a magnetic field) of this magnitude, then our microscope can readily resolve such a change. To extract, with statistical confidence, such a small change based on different cell populations is very challenging indeed and the sampling of ~100 cells quoted is on the optimistic side. This misunderstanding is repeated throughout the report of the replicate study.

*“However, we think that the major shortcoming of this study, that have led to the misinterpretation of the results [25], was making inference without directly comparing two effects (e*.*g. autofluorescence in the presence and in the absence of magnetic field). In fact, this is very common mistake leading to inappropriate analysis in the literature [50]. Making conclusion about an effect of some treatment should be based on a direct statistical comparison between a control and a treatment group [50]*.*”* This is the key point of contention, as addressed above. The great difficulty of looking for small magnetic effects in cells is the huge variability of the cells themselves. Each cell and region will have different concentrations and distributions of flavins. Therefore, to observe the very small effect of an applied magnetic field, needs averaging of a very large number of cells such that that the standard deviation drops below the size of the effect to be observed. This problem is completely solved by the realtime modulation of the magnetic field on a given region of a given cell during a single irradiation. In this case, the only changes observed are due to the magnetic field, whose strength and period can be controlled and correlation ensured. Thus this paragraph criticizes exactly what is beneficial about our approach. As our data indicate, we see a clearly correlated modulation of the fluorescence in all cells. The size of the modulation displays variability (see our supporting information^2^ fig. S1) but it is always observed. It is not observed in the reproduction due to the mismatched irradiation conditions.

“*In fact, in the study of Ikeya and Woodward [25] it is stated that “a large number of individual HeLa cells for different cell cultures prepared on different days” was used to analyze autofluorescence. Unfortunately, we could not find a definition of “a large number”, nor any other sample size determination*.” The reason we did not specify precisely the number of samples is that we are not performing any kind of statistical analysis. Each MFE measurement is for an individual cell and measuring more cells simply results in greater averaging and an increase in the signal to noise ratio as explained above. This is very clearly apparent in our experimental data. “However, only performing replicates is not enough; repeating experiments should be conducted in different days [53].” We repeated our measurement on many different days (more than three) over different weeks, but again, it is not relevant to our analysis, except to indicate that the observed effects were consistent, whenever the experiments were performed.

*“Another essential change we introduced, when analyzing the data for autofluorescence decay, was a background correction. It is absolutely necessary for accuracy and precision in quantitative fluorescence microscopy measurements, especially when quantifying weak signals [42, 47].”* This is not a change at all. All of our fluorescence data was also background corrected. Indeed in the reproduction, the authors focus a great deal on the measured ‘signal-to-noise’ ratio but what they actually present is the signal-to-background ratio. The critical signal-to-noise ratio that needs to be much smaller than the size of the MFE is a very different value. In our experiments, **Fig 3** shows the signal-to-noise ratio throughout the fluorescence decay. In addition, we report the background noise level in the absence of a magnetic field modulation in figure S7 in our paper^2^ which corresponds to a typical MFE-to-noise ratio of around 4 in a typical cell.

**Figure 3.**
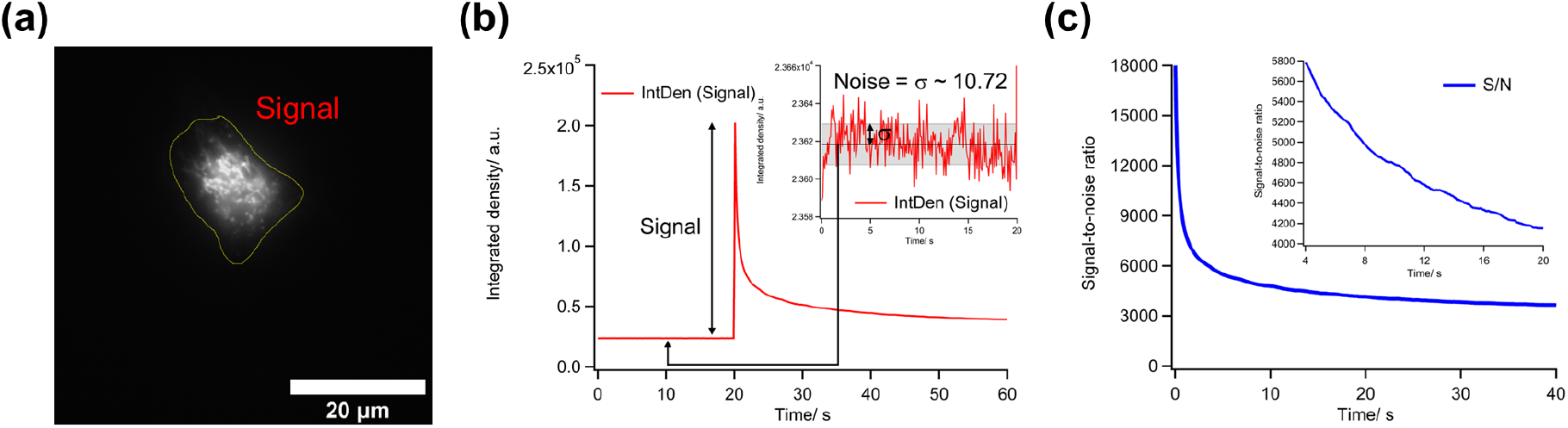
The signal-to-noise (SN) ratio of the autofluorescence of the Hela cell shown in the original study^2^. (a) The region of signal integration. (b)The integrated density change during the measurement. The zoom-in data in the inset shows camera noise. (c) The change of the SN ratio. The zoom-in data shows the change during a period used for the MFE analysis. All calculations of the integrated density were performed with the same method as the replicate study^1^.

*“We were puzzled by how the fractional MFE was estimated in the Ikeya and Woodward study [25]. The average autofluorescence decay curves were fitted using some exponential function [25]. We could not define; what kind of exponential function exactly was used. In the methods section “double exponential function” was stated, whereas in the main text and in the supplements “single exponential function” were mentioned [25]*.*”* There is no confusion over the fitting performed in our measurements. All the fluorescence decay curves were fitted with the following double exponential function for the baseline subtraction.

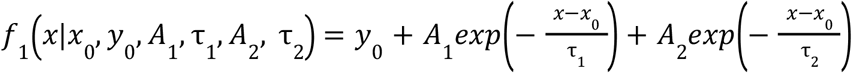

An equivalent fitting procedure is described in detail in a recent study^18^. In contrast, the single exponential fitting corresponds to an entirely different procedure on an entirely different data set. This single exponential fitting is used to measure the rate at which the fluorescence responds to square wave magnetic modulation - i.e. it determines the time response of the fluorescence to the application of a constant magnitude magnetic field^19^. The following single exponential function was used for the curve fitting.

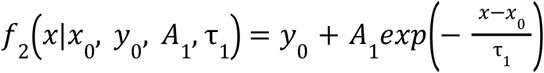

We think it is unreasonable to imply that our analysis method is unclear due to a misunderstanding of the use of two different fitting models for two different data sets.

### Data quality in the replicate study

Having argued that the reproduction is incapable of reproducing our experimental observations on theoretical grounds, based on the very different irradiation conditions, and then having defended our real-time magnetic field modulation based approach, we will conclude this rebuttal by highlighting significant problems with the experimental data presented in the reproduction study. Quoting again from that study, *“Some cells, even without applied field, show fluctuating “spikes” of autofluorescence during photobleaching course (Figures S8 and S9). In order to clearly examine these changes in autofluorescence, we sub-selected from 12 to 18 of the most “responding” cells (Figure 2A)*.*”* This is a clear problem with the measurement system / conditions in the reproduction study. In our original work, no such inexplicable spikes appear in the experimental data and for every individual cell, smooth fluorescence decays with low noise are observed. The spikes in the reproduction study are orders of magnitude larger than any possible magnetic field effects, and appear to be random in their timing. This significantly exacerbates the problem of statistical cell sampling, as each cell measurement has uncorrelated errors which once again requires substantial averaging to eliminate. It is not completely clear why these spikes are so prominent and of such great magnitude, but it seems likely that this is due to the high vertical resolution associated with the confocal microscope used in the reproduction. Any movement of e.g. mitochondria in the vertical plane would lead to them rapidly moving in and out of the vertical measurement slice and thus to sharp fluctuations in the observed fluorescence. Our measurements do not resolve vertically and in principle capture all the fluorescence in the region of interest over the whole vertical cell span, which completely eliminates such fluctuations. Again this was part of our original instrument design considerations, to alleviate as many sources of variability in the fluorescence signal as possible. The fluctuations are sufficient to essentially invalidate all the measurements in the reproduction study.

### Conclusions and recommendations

There are three important points made here that can be summarised as follows:

1. The spinning disc confocal scanning microscope used in the reproduction study renders the measurements incapable of suitably driving the photochemistry to allow for the observation of magnetic field effects on the ground state fluorescence.
2. The argument for statistical averaging across cells in field and no-field conditions is flawed as it is subject to the inherent cell and region variability and also the problem of non-uniform irradiation in the spinning disk arrangement (which means that the irradiation conditions may differ from cell to cell). Our original work which uses real-time magnetic field modulation eliminates the variability problem and the need for such statistical averaging.
3. The experimental data presented in the reproduction is incapable of resolving small changes in the fluorescence due to very large random fluorescence spikes. The origin of these spikes appears to be fluorescence structures moving in and out of the well-resolved vertical plane of fluorescence capture.

In many ways, the reproduction paper provides strong arguments for our original experimental approach. We would very much like to see the authors try again to reproduce our observations with appropriate irradiation conditions (i.e. removal of the scanning disc unit) and are confident that they will be able to reproduce our original observations. Having direct measurement tools to look at real-time changes in single cells is a necessary approach to be able to properly investigate small magnetic field induced changes at the cellular level.

We would like to strongly encourage other research groups with appropriate instrumentation to try to reproduce the results of our study. Taking into account the above discussions and speculating about other possible sources of problems, we would like to provide some simple recommendations for future reproduction studies:

1. The irradiation conditions are absolutely critical as the excitation light serves two roles - one for generating the fluorescence signal and the other for driving the magnetically sensitive photocycle. If the latter is not driven appropriately, magnetic field induced changes in radical concentrations will not be reflected in the observed fluorescence. We strongly recommend using an optical irradiation configuration that matches our own, as it is optimised for the photochemical reaction cycle.
2. In our study, cells were prepared in a particular way. In the first instance matching these conditions will increase the chance of observing the same responses. Because there are possibilities that endogenous flavin concentrations are dependent on cellular stress^20^ and cell confluence^21^. In the original work, to obtain consistent results, we precisely controlled the cell conditions before passaging into imaging chambers, the number of passaged cells, the culture medium volume and the incubation time before MFE measurements. The cell condition before passaging into the chambers was that two or three days of the log phase passed and the cell growth reached over 80% confluency. The number of the passaged cells was adjusted by diluting 150- or 200-fold from over 80% confluence on the surface of a 10 cm culture dish, so that the population density of the passaged cells on the surface of the chamber (0.95 × 0.95 cm) was the same for measurements on different days (judged by eye). The culture medium volume for the incubation used was 0.2 mL (giving 0.22 mL per 1 cm^2^ of bottom area) which follows the recommended medium volumes for cell culture. The incubation time before MFE measurements was 42 to 48 hours to allow the cells to recover from stress caused by the passage. After the incubation, we washed the chamber well twice with the PBS buffer to remove the medium completely (which causes high background signal as the medium contains riboflavin) and replaced it with the PBS buffer and measured the cells^2^.

